# Natural mutations of human *XDH* promote the nitrite (NO_2_^-^)-reductase capacity of xanthine oxidoreductase: a novel mechanism to promote redox health?

**DOI:** 10.1101/2023.03.24.533749

**Authors:** G. Massimo, R. S. Khambata, T. Chapman, K. Birchall, A. Shabbir, Nicki Dyson, K. Rathod, C. Borghi, A. Ahluwalia

## Abstract

Several rare genetic variations of human *XDH* have been shown to alter xanthine oxidoreductase (XOR) activity leading to impaired purine catabolism. However, XOR is a multi-functional enzyme that depending upon the environmental conditions also expresses oxidase activity leading to both O_2_ ^·-^ and H O and nitrite (·NO_2_ ^-^) reductase activity leading to NO. Since these products express important, and often diametrically opposite, biological activity consideration of the impact of XOR mutations in the context of each aspect of the biochemical activity of the enzyme is needed to determine the potential full impact of these variants. Herein, we show that known naturally occurring *hXDH* mutations do not have a uniform impact upon the biochemical activity of the enzyme in terms of uric acid (UA), reactive oxygen species (ROS) and nitric oxide (·NO) formation. We show that the His1221Arg mutant, in the presence of xanthine, increases UA, O_2_^·-^ and NO generation compared to the WT, whilst the Ile703Val increases UA and ·NO formation, but not O_2_^·-^. We speculate that this change in the balance of activity of the enzyme is likely to endow those carrying these mutations with a harmful or protective influence over health that may explain the current equipoise underlying the perceived importance of *XDH* mutations. We also suggest that targeting enzyme activity to enhance the NO_2_^-^-reductase profile in those carrying such mutations may provide novel therapeutic options, particularly in cardiovascular disease.

**Highlights:**

- Mutations of xanthine oxidoreductase modulate both its expression and activity
- The His1221Arg natural mutation increases xanthine oxidoreductase activity
- Raised xanthine oxidoreductase activity coupled with increased availability of nitrite substrate leads to increased NO provision

## Introduction

The pathophysiological role of xanthine oxidoreductase (XOR) (EC 1.17.3.2, xanthine:oxygen oxidoreductase), despite being discovered in 1902, remains incompletely understood. The gene (*XDH*) encoding for human XOR is transcribed as xanthine dehydrogenase (XDH), which is thought to be the main isoform expressed in humans under physiological conditions. However, under pathological conditions such as ischaemia-reperfusion, inflammation and hypertension XDH can be reversibly or irreversibly converted to the oxidase form of the enzyme (XO) via oxidation of cysteine residues (Cys535, Cys992) or proteolytic cleavage of lysine (Lys551, Lys569) respectively (1,2). XOR belongs to the molybdenum-containing enzyme family and exists as a homodimer of ∼300kDa with each monomer composed of three main domains: an 85kDa C-terminal domain containing the molybdo-pterin-binding site (Mo-Pt), an N-terminal domain of 20kDa containing two non-identical iron-sulphur (Fe-S) clusters, and an intermediate domain of 40kDa which is the flavin adenine dinucleotide (FAD) domain.

Both XDH and XO, at the Mo-Pt binding site, catalyse two consecutive hydroxylation reactions of hypoxanthine to xanthine and xanthine to the end product UA (3). However, differences exist between the two isoforms at the FAD site, where nicotinamide adenine dinucleotide (NAD^+^) is the main electron acceptor for XDH and is reduced to nicotinamide adenine dinucleotide hydrogen (NADH). In contrast, O_2_ is the only electron acceptor for XO, leading to the formation of superoxide anion (O_2_ ^-^) and hydrogen peroxide (H O) (4). It is noteworthy that under certain conditions such as ischaemia-reperfusion injury, characterised by increased levels of NADH, both XDH and XO operate as NADH-oxidase enzymes, with NADH oxidized at the FAD site and electrons used to generate O_2_ ^-^ and H O (5,6). This activity is independent of the Mo-Pt binding site but can be blocked by diphenyleneiodonium (DPI), a non-specific inhibitor of the flavin group (5,7).

Whilst XOR is important for healthy purine catabolism, elevation in activity of the enzyme and build-up of UA is pathogenic in gout (8) and in cardiovascular disease (CVD) progression (9), particularly in hypertension (10,11). Similarly, the production and accumulation of ROS, due to increased XOR activity, is implicated in numerous disease states characterised by an oxidative stress and endothelial dysfunction including hypertension (9). Numerous modifiable factors influence XOR expression and activity in disease including diet, cigarette smoke and alcohol consumption. In addition, non-modifiable factors exist and these include race, and ethnicity (12–16). These latter influencers intimate that genetics is an important factor to consider when assessing individual XOR activity and its functional impact. Indeed, naturally occurring mutations in h*XDH* have been identified, with two well-described nonsynonymous mutations leading to the rare disorders of hereditary Xanthinuria Type I and II. These mutations prevent purine metabolism causing consequent build-up of hypoxanthine and xanthine (17). In addition, some published polymorphisms are thought to be relevant for CVD and are localized to intron segments of the gene (18–20), in addition to nonsynonymous mutations (17,21–24). The minor allele frequencies (MAF) of these variants using the Genome Aggregation Database (https://gnomad.broadinstitute.org) and the International Genome Sample Resource (http://phase3browser.1000genomes.org/index.html) demonstrate a prevalence of 1-11%.

Recent large-scale studies interrogating genetic associations with gout and/or relevant CVD have not highlighted mutations in *hXDH*, but rather genes associated with UA metabolism and clearance (25–27). However, some relatively small studies have linked specific h*XDH* variants with common disease, particularly in Asians. For instance in 196 Japanese individuals, a single nucleotide polymorphism (SNP) in the promoter region of h*XDH*, when reproduced in vitro demonstrated altered XOR expression levels (28). Another small study (185 patients, 370 controls) described specific variants linked with oxidative stress and hypertension (20) and resonates with a study in Europeans with hypertension linking UA levels and *hXDH* mutations (18). Many of the identified variants have been tested for functional impact upon enzyme activity in vitro. Kudo et al. in 2007, found two SNPs, 3662 A>G and 2107 A>G, leading to His1221Arg and Ile703Val mutations respectively, showing nearly a two-fold higher UA synthesis when compared with WT (22). However, how these variants impact all aspects of XOR activity particularly the NO_2_^-^ reductase activity of the enzyme either alone or in context with its other biochemical activities is unknown(29). To explore this, we expressed several well described h*XDH* mutations in HEK- transfected cells and assessed all three aspects of XOR activity to better ascertain any potential functional impact of these mutations.

## Material and Methods

### hXDH mutation computational predicted crystal structure

Using the 2EIQ structure, which is a 2.6 Å resolution full length human structure we analysed the impact of 4 very well described hXDH mutations upon the structure of the protein. The Schrodinger protein preparation wizard was used to assign bond orders, add hydrogen atoms and optimise tautomer, rotamer and protonation states at pH 7.0, finishing with a restrained minimisation (OPLS2005 force field with convergence of heavy atoms to 0.3Å RMSD) to relieve any crystallographic strain. The amino acid mutations were performed in Maestro (2019 version) by selecting the lowest energy rotamer, then performing a simple minimisation restricted to residues within 5Å of the mutated residue.

### *hXDH* Subcloning

h*XDH* wild type (WT) cDNA (kind gift of Professor Richard M. Wright) was excised from a pcDNA3.1-myc-hisA plasmid vector using NheI-HF and EcoRV-HF (New England Biolab) restriction enzymes (REs). Following purification, a Kpn-I sequence at the 3’ of the cDNA was introduced and then cut with NheI-HF and KpnI-HF and a GFP-tagged ligation product, pEGFP-N1 – h*XDH* introduced for visualisation of expression. Sequence was verified via Sanger sequencing (Applied Biosystems 3730 capillary sequencer) and interpreted using BioEdit Sequence Alignment Editor Software before proceeding with the site directed mutagenesis.

### Site directed mutagenesis

Mutagenic primers, shown in Table S1 (This article contains supporting information), were designed using NEBaseChanger software (http://nebasechanger.neb.com). Using a Q5^®^ Site-Directed Mutagenesis Kit (New England Biolab) we reproduced 4 non-synonymous mutations (Table 1). These variants were chosen due to previous studies identifying substantial change in conventional XOR activity and include mutations that lead to direct alterations in structure to the Mo-Pt (Ile703Val, His1221Arg, Asn909Lys) and FAD (Trp336Ala/Phe337Leu) subunits of the enzyme(22,30).

**Table 1:** Summary table indicating nucleotide substitution, exon position, amino acid changes and SNP ID for each investigated hXDH mutant.

| Variant | Position | Nucleotide Change | Mutation | Exon | Subunit position | SNP ID | MAF | MAF (gnomAD exomes r2.1.1) |  |  |  |
| --- | --- | --- | --- | --- | --- | --- | --- | --- | --- | --- | --- |
|  |  |  |  |  |  |  |  | European | Asian | American | African |
| Asn909Lys | 2727 | C > A | SNP | 25 | Mo-Pt | rs566362 | <0.01 |  |  |  |  |
| His1221Arg | 3662 | A > G | SNP | 34 | Mo-Pt | rs376191022 | <0.01 |  |  |  |  |
| Ile703Val | 2107 | A > G | SNP | 20 | Mo-Pt | rs17011368 | 0.05 | 0.032 | 0.0187 | 0.02 | 0.011 |
| Trp336Ala/<br>Phe337Leu | 1006/1007<br>1009 | TG > GC<br>T > C | Artificially produced | 11 | Rely System |  |  |  |  |  |  |

PCR conditions used for each mutant isoform are shown in Table S2 (This article contains supporting information). NEB 5-alpha competent *E. coli* cells (New England Biolab) were used for bacterial transformation. Up to three resistant colonies per mutation were selected and plasmid DNA was extracted using a QIAprep Spin Miniprep Kit (Qiagen) and quantified using a nanodrop spectrophotometer (ND.1000). Mutations were confirmed via Sanger Sequencing as above using primers reported in Table S3 (This article contains supporting information). Mutated Plasmid cDNAs were then prepared and extracted using the Qiagen^®^ Plasmid maxi Kit.

### HEK-293 transfection and stable cell line formation

Human embryonic kidney cells (HEK-293) were chosen to stably express the wild-type and all mutated variants as previous findings suggest that they do not express XDH (31) an observation confirmed in The Human Protein Atlas (https://www.proteinatlas.org/ENSG00000158125-XDH/cell+line). To produce stable cell lines of WT and mutated hXDH, 3 x 10^5^ cells per well were seeded into a 6-well plate in high-glucose Dulbecco’s modified Eagle’s (DMEM) medium (Merck, UK) enriched with 10% FBS (Sigma-Aldrich), 1% l-glutamine (Sigma-Aldrich), 1% penicillin/streptomycin, until at 60-70% confluency. The plasmid constructs were transfected using Lipofectamine® 2000 Reagent (Invitrogen) as per the manufacturer’s instructions72h after transfection 1.2mg/ml of G-418 (Sigma-Aldrich), previously determined via a kill-curve, was added to DMEM normal medium until single colonies appearance. 4-5 colonies per mutations were then expanded and XOR expression levels checked via immunoblotting. Only colonies with the highest expression profile were expanded, amplified, and stored in liquid nitrogen for future experiments.

### Cell culture and cell lysate extraction

HEK-293 cells were maintained in high-glucose DMEM medium supplied with 10% FBS, 1% l-glutamine and 1% Penicillin/Streptomycin, and G-418 (Sigma-Aldrich) 0.3mg/ml. Cells were cultured under 5% CO_2_ and at 37 °C and split every 72h at 70-80% confluency. To prepare cell lysate, cells were cultured till 80-90% confluency and cell pellets stored at -80 °C until the day of use. For protein extraction in 200μl PBS containing 1% Triton X, with 5.7μM benzamidine, 1.5μM antipain, 0.15μM aprotinin, 4.2μM leupeptin, 1.5μM pepstatin A and 400μM AEBSF protease inhibitor was added to the cell pellet and the pellet mechanically disrupted using a 25G needle (0.5mm). The resulting homogenate was centrifuged at 4**°**C for 10 mins at 14,000 RPM and the supernatant collected. Protein concentration was determined using a Pierce**®** BCA Protein Assay Kit (Thermoscientific, UK) according to manufacturer guidelines.

### Quantitative real-time PCR

mRNA expression was determined in HEK-293 stable cell lines using RT-PCR. The primer sequences used are shown in Table S4 (This article contains supporting information). 5 x 10^6^ cells were processed as per the manufacturer’s instruction using the NucleoSpin RNA, Mini kit for RNA purification (Macherey Nagel, MN). RNA was converted to cDNA and RT-PCR assays conducted using SYBR green ROX mix (Thermo Scientific Abgene, UK) in 384 well plates. An ABI7900 HT Realtime PCR System (Applied Biosystems**^®^,** Life Technologies, UK), with SDS 2.3 computer software, was used to run and analyse plates. Each sample was measured in triplicate and the average taken to represent n=1. Gene expression was measured relative to β-actin and GAPDH housekeeping genes, and expressed relative to h*XDH* WT.

### Immunoblotting

Cell lysates were subjected to SDS/PAGE (0.1% w/v) immunoblotting analysis using an anti-human XOR rabbit antibody (1:2000, Abcam 133268 RRID: AB_11154903). Briefly, 50μg of protein of each cell extract, was prepared for electrophoresis and loaded onto an 8-16% Mini-Protean TGX gels (Bio-rad). Following this, electrophoresed proteins were transferred via a semi-dry transfer method to a 0.2µm nitrocellulose membrane (Amersham™ Proton™). Red Ponceau solution was used to confirm transfer prior to overnight incubation with primary antibody. The membrane was then washed and then incubated with an anti-rabbit secondary antibody (1:5000, InvitrogenAB_2536381) for 1 hour. The levels of protein were expressed relative to GAPDH expression (1:5,000, ThermoFisherScientific #AM4300, RRID: AB_2536381). The nitrocellulose membrane was exposed to a 1:1 v/v solution of Clarity Western ECL Substrate (Bio-rad) for 5 mins and chemiluminescence quantified and analysed using FluorChem E software (ProteinSimple).

### Confocal Microscopy

A 1% gelatin (Sigma-Aldrich) pre-coated 12mm diameter coverslip (Epredia^TM^ Menzel^TM^, Fisher Scientific) was placed in each well of a 24-well plate and 10^5^ cells added and cultured until 70-80% confluency. Cells were then fixed with 4% paraformaldehyde for 15 minutes at room temperature after which wells were rinsed three times with PBS prior to addition of 500μl permeabilization solution (PBS + 0.25% Triton X-100). After washing with PBS a 2.5% BSA solution was applied for 1 hour and then incubated with primary anti-human XOR rabbit antibody (1:2000, Abcam 133268 RRID: AB_11154903) and left over-night at 4 °C. The following day coverslips were washed with PBS and incubated with the secondary antibody, Alexa Fluor^TM^ 555 donkey anti-rabbit (1:250, Life Technologies, RRID: AB_10892947) for 1h at room temperature. Each coverslip was then washed prior to incubation with PBS + DAPI (1:5000) staining solution. For each mutant two coverslips were analysed for each experiment with XOR expression visualized under a confocal microscope (ZEISS LSM 880 with Airyrscan) and images analysed with ImageJ software.

### Flow cytometry for XOR

5x10^5^ cells were fixed (Intracellular fixation and permeabilisation buffer, Thermofisher, UK), the cells permeabilised (Intracellular fixation and permeabilisation buffer, Thermofisher, UK) and then incubated with 2% donkey serum followed by anti-human xanthine oxidase (1:1000; ab133268, Abcam, UK; RRID: AB_11154903), for 30min at 21 °C. Cells were then washed and the secondary antibody (donkey anti-rabbit IgG preadsorbed, Abcam, RRID: AB_2715515) incubated for 30min 21°C. Samples were analysed using a BD LSRFortessa Flow Cytometer, and data recorded using BD FACSDiva analysis software. Histogram images were produced using FlowJo v10.7.2.

### Pterin-based fluorimetric assay of XDH/XO activity

To determine the relative proportions of XDH versus XO activity for each mutant we used a modified protocol as described by Beckman et al. (1989)(32). 1000μg of protein, obtained as above, was mixed with pre-warmed PBS (50mM) + Na_2_-EDTA (0.1 mM) buffer pH 7.4 in a final volume of 200μl. Baseline fluorescence was measured every 20 sec for 5 min. XO activity was measured by adding pterin (10μM) whereas XDH + XO activity was quantified in the presence of 10μM methylene blue. The reaction was stopped by adding 10μM allopurinol before reading the fluorescence produced by isoxanthopterin (1μM), used as an internal standard. Stock solutions were made fresh every day; 100mM of pterin and 10mM of isoxanthopterin stock solutions were prepared in NaOH 1N, and then serially diluted in PBS buffer. 10mM methylene blue was prepared in MilliQ H_2_O, and diluted as required. Fluorescence was measured using a Tecan i-control plate reader (infinite M200 Pro) excited at 345nm and with emission at 390nm. The fluorescence data was corrected for the immediate fluorescence increase after isoxanthopterin addition, and calculated using the following formula:

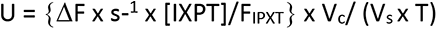

where U is the enzyme activity in μmol s^-1^ g protein^-1^, ΔF is the change per second in fluorescence intensity obtained for pterin or methylene blue, [IXPT] is the concentration of the isoxanthopterin (µM) added at the end of the assay. F_IPXT_ is the immediate increase of fluorescence obtained after isoxanthopterin addition. V_c_ is the final volume used in mL, V_s_ is the volume of sample added in mL, and T indicate the concentration of the homogenate (mg ml^-1^).

### Superoxide quantification via LECL

For O_2_^·-^ quantification via LECL 100μg protein was added to each well in a 96-well plate, solid white with a flat and transparent bottom (Greiner Bio-One^TM^). Briefly, red phenol-free Hank’s Balanced Salt Solution (HBSS, Lonza) supplemented with HEPES (1mM) pH 7.4 was mixed in a 1:1 v/v ratio with Na_2_CO_3_/NaHCO_3_ 0.1M (4:1 v/v) buffer, pH 11.0; final pH was adjusted with NaOH or HCl 1M to 10.4. In some cases, protein lysates were incubated with pharmacological inhibitors, febuxostat (1μM) or DPI (10μM) for 30 min at 37°C. Experimental substrate(s) xanthine, NADPH or NADH were added at 100μM per well, followed by 10μM lucigenin (Sigma Aldrich, UK) as previous studies (33). Luminescence was immediately analysed using a Perkin Elmer Wallac 1420 Victor2 plate reader (Perkin Elmer, USA), O_2_^·-^ anion production was read for 50 repeat readings with 30 sec interval between each repeat. Each sample was run in duplicate, and the results averaged to give an n=1.

### UA quantification

1.3 x 10^6^ cells were seeded into 75cm^2^ flasks. After 72h the medium was replaced and supplemented with xanthine 20μM or 100μM and incubated at 37 °C for 3h, after which cell pellets were collected as above, and stored at -80 °C for future assessment. One half was used for protein extraction and the other half used for UA quantification using a UA Assay Kit (MAK077, Sigma-Aldrich). Cells were disrupted mechanically, and the sample centrifuged at 15,871 RCF at 4 °C. The supernatant was then added to a 96-well plate for colorimetric quantification using a spectrophotometer at 570nm. UA values were obtained by interpolation from a standard curve and then normalized to the protein concentration.

### NO_2_^-^-reductase assessment

NO_2_^-^-reductase activity was measured in cell lysate using gas-phase chemiluminescence as previously described (34). Cell pellets were homogenised with homogenization buffer devoid of Triton X-100. Experiments were performed in a sealed 10 ml glass reaction chamber containing citric acid/Na₂HPO₄ buffer at pH 6.8 (representing acidosis), and NaNO_2_ (10-1000 μM) in a total volume of 1 ml. This solution was bubbled with nitrogen gas (100%) via an ·NO scrubbing air filter (Sievers, USA) to eliminate all O_2_. Headspace ·NO concentration was measured in parts per billion by continuous sampling using a 280A ·NO Analyzer (Sievers, USA). The impact of biological tissue on NO production from NO_2_^-^ was determined by the addition of 250μg of protein and measurement of chemiluminescent signal of·NO over 2 min, calculating the rate of ·NO production (nmol g protein^-1^ s^-1^) from the area under the curve (AUC). Involvement of XOR was confirmed by assessment of the effect of pre-treatment with febuxostat (10µM) or vehicle control (0.05% DMSO) for 30 min. Furthermore, to establish whether non-synonymous mutations influenced the enzyme selectivity toward Mo-Pt or FAD reducing agents and consequently the NO_2_^-^-reductase activity, nitrite-reductase activity was carried out in presence of xanthine (20µM) and NADH (100µM).

### Statistical analysis

Statistical analysis was conducted using GraphPad Prism 9 software. All data are expressed as mean ± SEM. For statistical comparison, either an unpaired student t test analysis or one-way ANOVA with Dunnett’s post hoc analysis, or two-way ANOVA followed by Dunnett or Sidak’s post hoc analysis for multiple group comparison were used as appropriate. Any *P* value less than 0.05 was used to infer significance. For all datasets an n=7 was achieved. Where the n value is less than this this is due to technical failure. The n value of 7 was set according to our preliminary experimental findings with assessment of superoxide levels in control untransfected cells and those transfected with our preliminary data with 5 independent experiments (separate cultures conducted on different days of the source cells) and demonstrated that cell homogenate incubated with Xanthine (100μM) gave an average count per second (CPS) of 146 (SD=14) and *hXDH* transfected cells gave 440 (SD=38). Thus, with an α=0.05 and a power of 95% an n=5 independent experiments are required to observe statistical differences. To account for potential technical failure we increased this number to 7.

## Results

### Computational analysis of 3D structure of mutated XDH

To explore the impact of *hXDH* mutations upon enzyme function we first assessed four well described *hXDH* mutations upon the predicted structure of the protein; namely Trp336Ala/Phe337Leu, Ile703Val, His1221Arg and Asn909Lys (Table 1, Figure 1)(22,30). Importantly, these mutations occur naturally within the human population except for the Trp336Ala/Phe337Leu, which was created experimentally and selected specifically due to its localisation at the FAD site of the enzyme (30).

**Figure 1.**
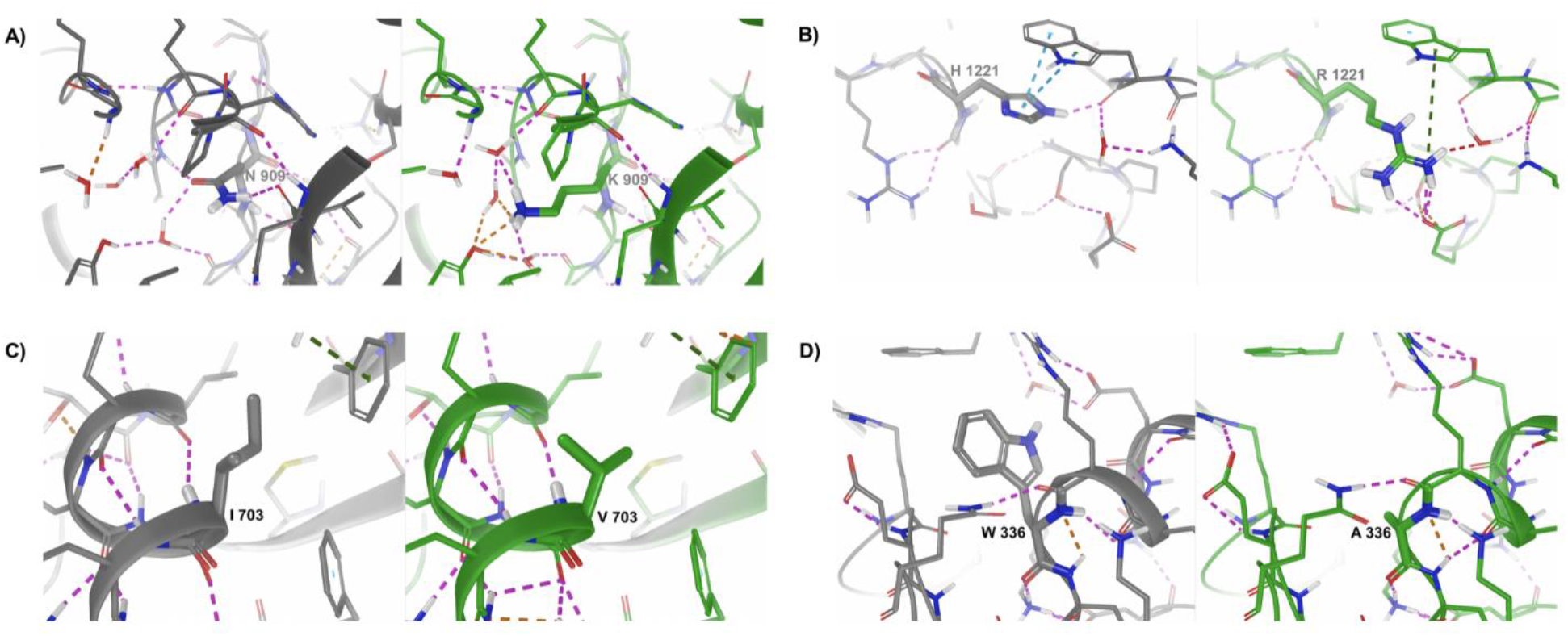
Impact of *hXDH* mutations on XOR protein structure. Wild type structure (2E1Q) is shown in grey beside the mutated structure shown in green. Residues within 5Å of the mutated residue are shown as thin sticks, whilst the mutated residue is shown as thick sticks. Interactions are indicated by dashed lines, with H-bonds in magenta, aromatic in cyan, cation-pi in dark green, with other non-bonded short contacts shown in orange or red according to increasing severity. A) N909K, B) H1221R, C) I703V and D) W336A

The 2EIQ structure, which is a 2.6Å resolution of the full-length human structure already contains a mutation (E803V) at the Mo-Pt site which is known to affect function, however this does not impact the structure in any way based on comparison with 2CKJ. Residues 164-191 are not visible in the density, nor are they in the other human structure 2CKJ. Given that this loop is exposed, long and flexible any predicted conformation would be of low reliability, so we did not pursue to build this part of the protein. Figure 1 shows the predicted structures of the protein in presence of the mutations assessed. N909 is largely buried, but not too tightly enclosed. Interactions between its sidechain and the peptide backbone are lost on mutation to lysine and adjustment of the solvent network observed in the crystal is also required to accommodate the change. H1221 interacts with the backbone and sidechain of W1329, likely stabilising the loop structures in this region. Mutation to arginine is a substantial change and whilst the aromatic interactions with W1329 are lost, there are still some interactions due to the cationic nature of the arginine, which also engages with E1217. I703 is engaged in hydrophobic interactions with F865, F708 and I699, helping to stabilise packing of the helix against the beta sheet. Mutation to valine is quite conservative but reduces hydrophobic contact. W336 is tightly enclosed and loss of packing interactions on mutation to alanine is expected to destabilise interactions in this region possibly resulting in conformational perturbations beyond those reflected in the modelled structure.

### Characterization of hXDH WT and mutant cell lines: Sanger sequencing chromatograms, mRNA quantification and XOR expression level

Sanger sequencing confirmed WT and mutant sequences as expected (Figure 2A-D). qPCR analysis revealed a significant difference of XDH mRNA and protein expression levels across the cell lines, with WT showing an mRNA level 2.0-fold higher than Trp336Ala/Phe337Leu, 2.5-fold when compared to Ile703Val, 3.4-fold higher than His1221Arg, and 5.0-fold when compared to Asn909Lys (P<0.0001, Figure 2E). Immunoblotting revealed the presence of two bands, as described previously, attributed to XDH and XO (35,36), at ∼150 and ∼100 kDa respectively. For quantification the band for the intact protein at 150kDa was used, although inclusion of the second band does not alter the pattern of expression. The analysis confirmed a significantly higher expression of XOR protein of the WT compared to the variants. Interestingly, for Trp336Ala/Phe337Leu we observed a lower level of protein than mRNA detection intimating perhaps that the mutation exerts a negative impact on the maturation or stability of the protein. In contrast, for Asn909Lys the opposite effect was observed (Figure 2F). The protein expression levels were confirmed by confocal microscopy (Figure 2G) and flow cytometry (Figure S1). Magnification of the images from the confocal assessments identified similar localisation of the WT and mutant proteins largely within the cytoplasm (Figure 2G). Total protein concentrations of collected cell lysates across the distinct cell lines were similar, except for Trp336Ala/Phe337Leu (23.05µg/µl) which produced higher levels than HEK cells alone (17.44 µg/µl).

**Figure 2.**
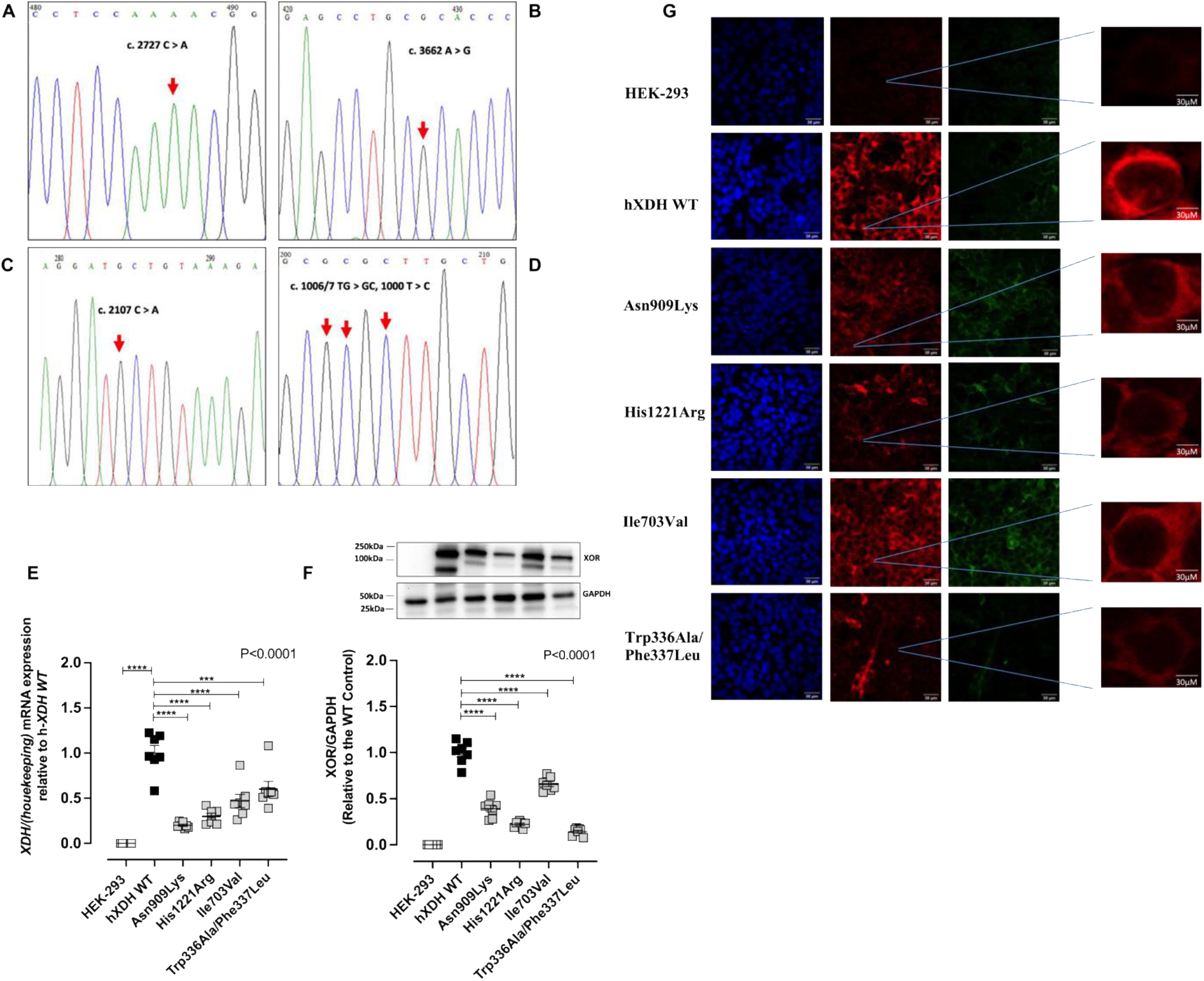
Site specific mutations of h*XDH* impact cellular expression levels. Sanger sequencing chromatograms of site-specific mutation sourced in the pEGFP-N1 plasmid vector are shown for A) 2727 C>A leading to Asn909Lys mutation, B) 3662 A>G accounting for His1221Arg, C) 2107 A>G leading to Ile703Val, D) triple mutation 1006/7 TG>GC + 1009 T>C leading to the Trp336Ala/Phe337Leu. E) shows mRNA levels of the various mutants versus the WT and F) representative immunoblotting and its quantification of WT and mutated hXOR. G) shows XOR expression detected via immunocytochemistry with DAPI (blue), XOR (red) and GFP (green). Magnification X 40; scale bar 30µm. All data are expressed as mean ± SEM of data and compared using one-way ANOVA with Dunnett post hoc analysis comparing to the hXDH WT control. Post- test significance shown as ***p<0.001, ****p<0.0001.

Since the mutations exerted a differential impact upon XOR protein expression, comparisons between mutants were made both before and after normalization to XOR expression levels in the same lysate determined using 50µg of cell lysate for immunoblotting detection. This amount of cell lysate protein was used for this since it provided an indication of XOR within the linear phase of expression (Figure S2A and B).

### Genetic variants of hXDH influence the XO:XDH activity relationship

The proportions of XDH (54%) and XO (46%) activity of the WT relative to total pterin oxidation activity (0.2389 ± 0.01 nmol/g/s, n=7) revealed statistically significant differences in the oxidative and dehydrogenase activities of the distinct mutations (Figure 3 B-D). In untransfected HEK-293 cells, total XDH+XO activity was negligible (0.0012 ± 0.0008 nmol/g/s, n=6). The Asn909Lys (XDH+XO=-0.0005 ± 0.0004 nmol/g/s) and Trp336Ala/Phe337Leu (-0.00008 ± 0.0009 nmol/g/s) mutations eliminated all activity of the enzyme in this assay. The non-synonymous mutation His1221Arg showed a similar total activity as that of h*XDH* WT, however the proportion of XO activity was substantially greater than that seen in the WT (∼70%, Figure 3E) suggesting a potential pro-oxidative effect of the mutation. When normalised to total XO/XDH activity the level of XO activity for Ile703Val was also statistically significantly higher than the XDH proportions (Figure 3D and E).

**Figure 3.**
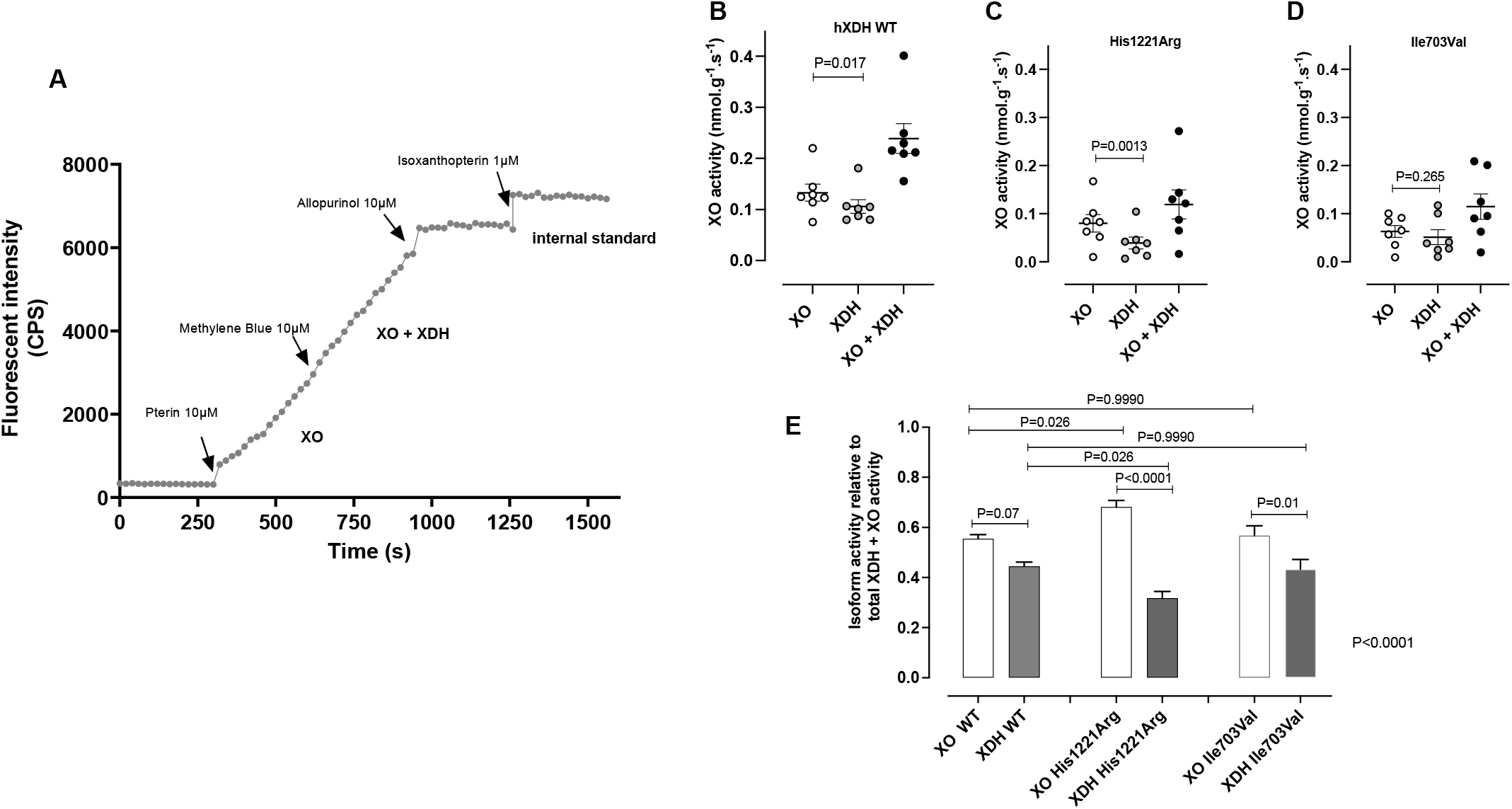
Genetic mutations influence XDH conversion to XO. Panel A) shows a representative trace of XO and XDH+XO fluorometric activity in the presence of pterin and methylene blue respectively over 300s intervals of recording. Panels (B-D) show the absolute activity of XO, XDH, and XO + XDH in hXDH WT, His1221Arg, and Ile703Val. Figure E) shows the XO and XDH activity relative to total XO + XDH activity (n=6). Pterin, methylene blue, allopurinol were used at a final concentration of 10µM, whereas isoxanthopterin, the internal standard, was used at 1µM final concentration. Fluorescence was recorded every 20 seconds for 5 minutes. Data are expressed as mean ± SEM of data and compared using a paired t-test comparing XO and XDH activity for B-D. For E statistical significance was determined using one way ANOVA (P value in right hand corner) with Sidaks post-test comparing XO and XDH activity for each genotype and XO WT versus XO activity of each mutation and XDH WT versus XDH of each mutation, with adjusted post-test significance accounting for multiple comparisons shown.

### ROS production is influenced by genetic variations

The observations above are further supported by our findings utilising the lucigenin chemiluminesence-based method of O_2_^·-^ measurement. His1221Arg and Ile703Val, statistically significantly increased O_2_ reduction compared to HEK-293. On the contrary, the C-terminal non-synonymous mutation, Asn909Lys, along with the artificially produced double mutation Trp336Ala/Phe337Leu, showed no residual oxidase activity implying a plausible conformational alteration of the tertiary structure around the Mo-Pt site (Figure 4). Confirmation of XOR and FAD site involvement in xanthine-driven O_2_^·-^ generation in this assay, was demonstrated by block of the response with febuxostat and DPI respectively (Figure 4B and C). As expected, whilst both inhibitors blocked xanthine driven O_2_^·-^ generation in WT, only DPI blocked NADH or NADPH driven radical generation. Of note, febuxostat caused an elevation of O_2_^·-^ generation in the presence of both these substrates although this did not reach statistical significance. To confirm that XOR overexpression did not influence NADPH oxidase expression and activity we also exposed our cell lines to NADPH. As expected, NADPH oxidase activity was not affected by the overexpression of XOR with the O_2_^·-^ quantification consistent throughout the mutations (Figure 4F). Since the mutations exerted a differential impact upon protein expression, O_2_^·-^ measurements were normalised to XOR expression level estimated from the same lysates using Western blotting analyses. This normalisation demonstrated that His1221Arg mutation resulted in a significantly higher O_2_^·-^ production compared to hXDH WT in the presence of either xanthine or NADH (Figure 4G and H) whilst the other mutations showed no statistical difference in oxidase activity to h*XDH* WT in the presence of xanthine. Although most trended to be lower particularly Asn909Lys and Trp336Ala/Phe337Leu (p=0.07) and, in the presence of NADH, Trp336Ala/Phe337Leu and His1221Arg showed increased activity relative to WT (Figure 4H).

**Figure 4.**
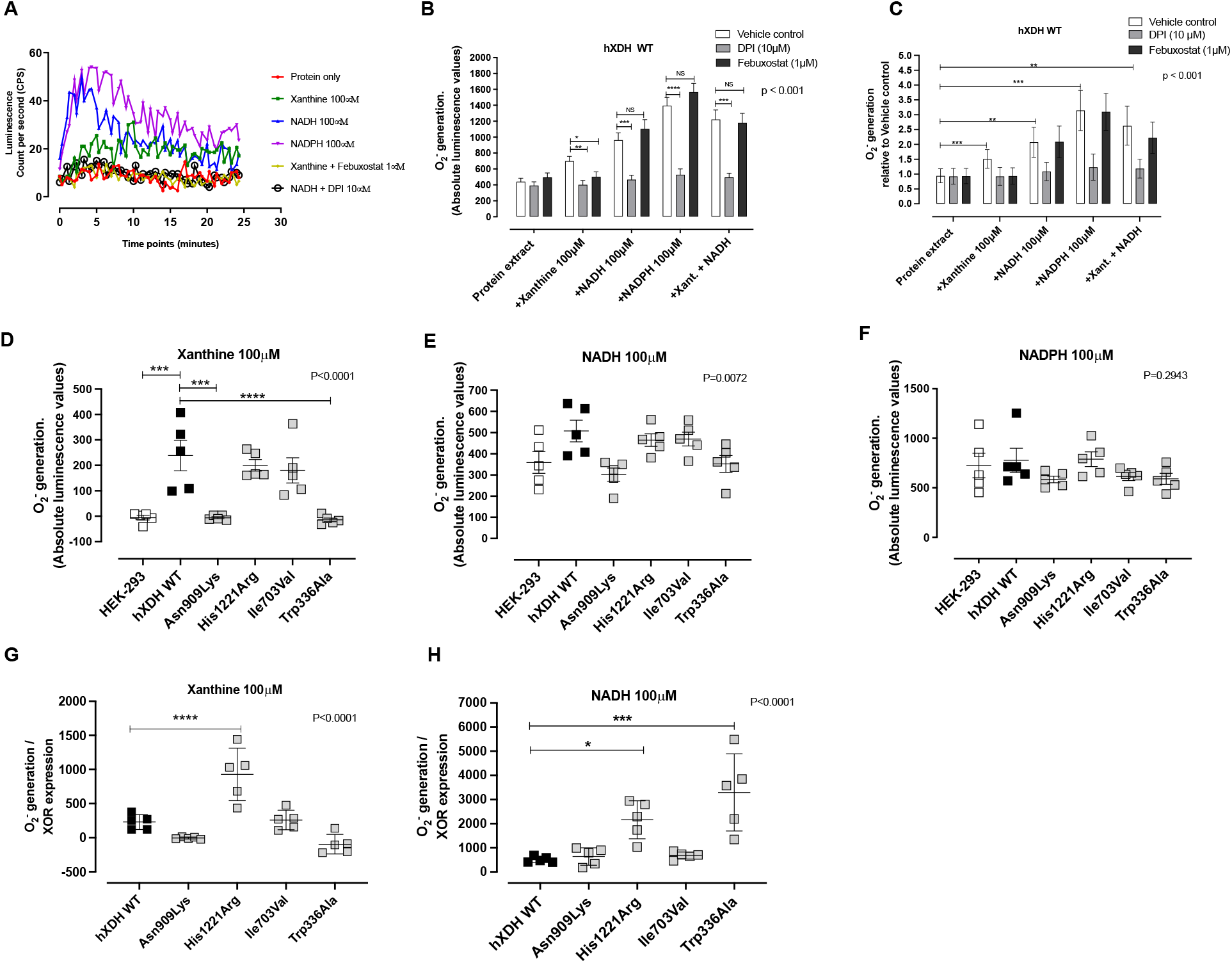
Impact of *hXDH* mutations upon superoxide anion (O_2_^-^) generation. Panel (A) shows representative traces of O_2_^·-^ quantification over 25 min of reading via lucigenin-enhanced chemiluminescence reaction (LECL) in the presence of 100µg of cell lysate of WT and mutated *hXDH* cell lines in the absence and presence of Mo-Pt-reducing substrate Xanthine (100µM), FAD- reducing substrate NADH (100µM), and NADPH oxidase-specific substrate NADPH (100µM). Panels (B-C) show O_2_^·-^ generation over 25 min in the absence and presence of a Mo-specific inhibitor, febuxostat (1µM), and FAD inhibitor diphenyleniodonium (10µM) in HEK-293 and *hXDH* WT stable cell lines respectively. Panel (D-F) show O_2_^·-^ quantification in presence of xanthine (D), NADH (E) and NADPH (F). Finally O_2_^·-^ quantification normalised to XOR expression level are shown for the cell lines in the presence of xanthine (G) and NADH (H). Data are presented as mean ± SEM. Statistical difference between groups was measured using two-way ANOVA for B and C with post-tests comparing each vehicle control response to the response in either DPI or febuxostat treated samples. For D-H one-way ANOVA followed by Dunnett’s post hoc analysis comparing to the hXDH WT control was conducted with post-test significance shown as *for p<0.05, *** for P<0.001 and **** for P<0.0001

### Differences in the genetic sequence alter UA production either in presence or absence of xanthine

To explore the impact of the mutations upon activity of the enzyme at the Mo-Co site we measured UA synthesis. As expected, baseline levels of UA were elevated in hXDH WT versus untransfected HEK-293 controls (Figure 5A). His1221Arg and, in particular, Ile703Val exhibited significantly increased levels of UA when compared to untransfected controls, with Ile703Val resulting in higher levels than hXDH WT (Figure 5A). To compare enzyme activity between variants more closely we normalized UA values to XOR expression levels. This revealed that only the His1221Arg mutation, in line with previous observations, was associated with a statistically significant raised urate level compared to hXDH WT (Figure 5B). Surprisingly, despite elimination of all O ^-^ generating activity UA levels were unchanged compared to WT with the Asn909Lys mutation. These results suggest that this non-synonymous substitution did not alter the active site cavity at the Mo-Co domain in a manner that prevented access and binding of xanthine. The double mutation isoform Trp336Ala/Phe337Leu increased UA levels compared to the WT but this did not achieve statistical significance (Figure 5B). Exposure of each cell line to xanthine demonstrated concentration dependent elevations for all mutations whilst untransfected cells did not respond (Figure 5C-H).

**Figure 5.**
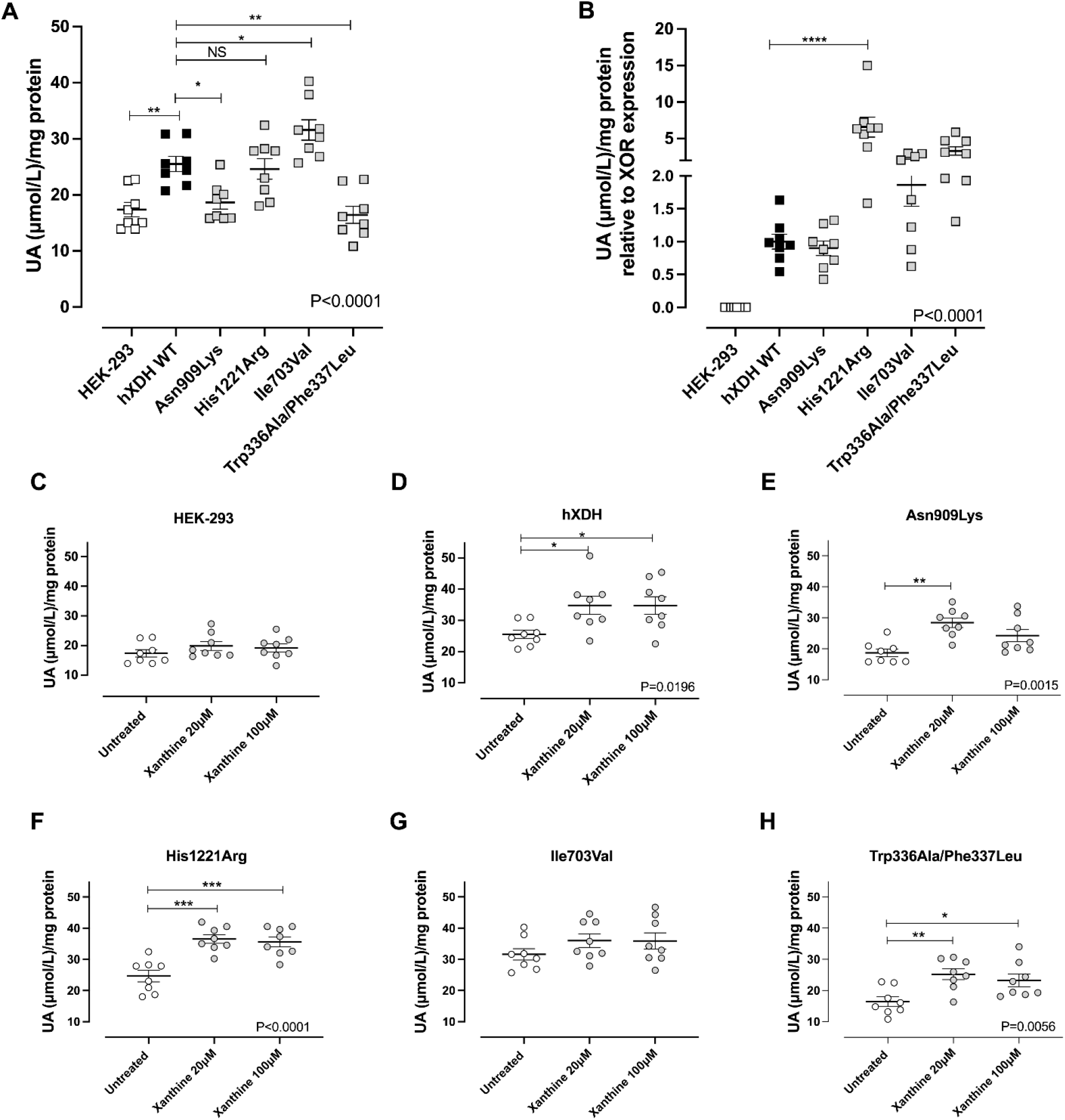
Mutations in XOR influence uric acid (UA) production. Panel (A) shows UA levels in the XDH WT and mutant cell lines at baseline relative to total protein concentration or following normalization to XOR expression (B) determined by Western blotting of the same sample. Panels C-H show the absolute UA levels generated from 4 x 10^6^ million cells incubated with xanthine 20- 100µM. Data are shown as mean ± SEM. Statistical analysis was conducted using one-way ANOVA with Dunnett’s post-hoc analysis for comparison to hXDH WT of n=8 independent experiments.

### *Improved NO_2_^-^* reductase activity of specific h*XDH* mutants

We next explored whether specific hXDH mutant forms might influence the NO_2_^-^-reductase capacity and so the generation of the beneficial molecule ·NO. The expression of hXDH WT in HEK-293 led to an increase in NO_2_^-^ -reductase activity above that with untransfected HEK-293 control. This elevation was also evident with both the His1221Arg and Ile703Val mutants (Figure 6A). Interestingly, both mutants Asn909Lys and Trp336Ala/Phe337Leu did not appear to show any differences in terms of absolute ·NO formation in comparison to untransfected cells (Figure 6A). When the NO_2_ ^-^-reductase activity was normalised to XOR expression level, His1221Arg exhibited an overall statistically significant higher NO_2_ ^-^-reductase activity compared with the hXDH WT (Figure 6B,) with a slight trend for elevation with Asn909Lys and also Ile703Val at baseline and in the presence of reducing substrate confirming a general trend for increase with all of the mutations despite clear differences in the O_2_^·-^ generating activities of these mutations (Figure 6C and D). Interestingly, whilst febuxostat prevented the activity for most mutations it did not impact activity in cells expressing Asn909Lys and Trp336Ala/Phe337Leu (Figure 7).

**Figure 6.**
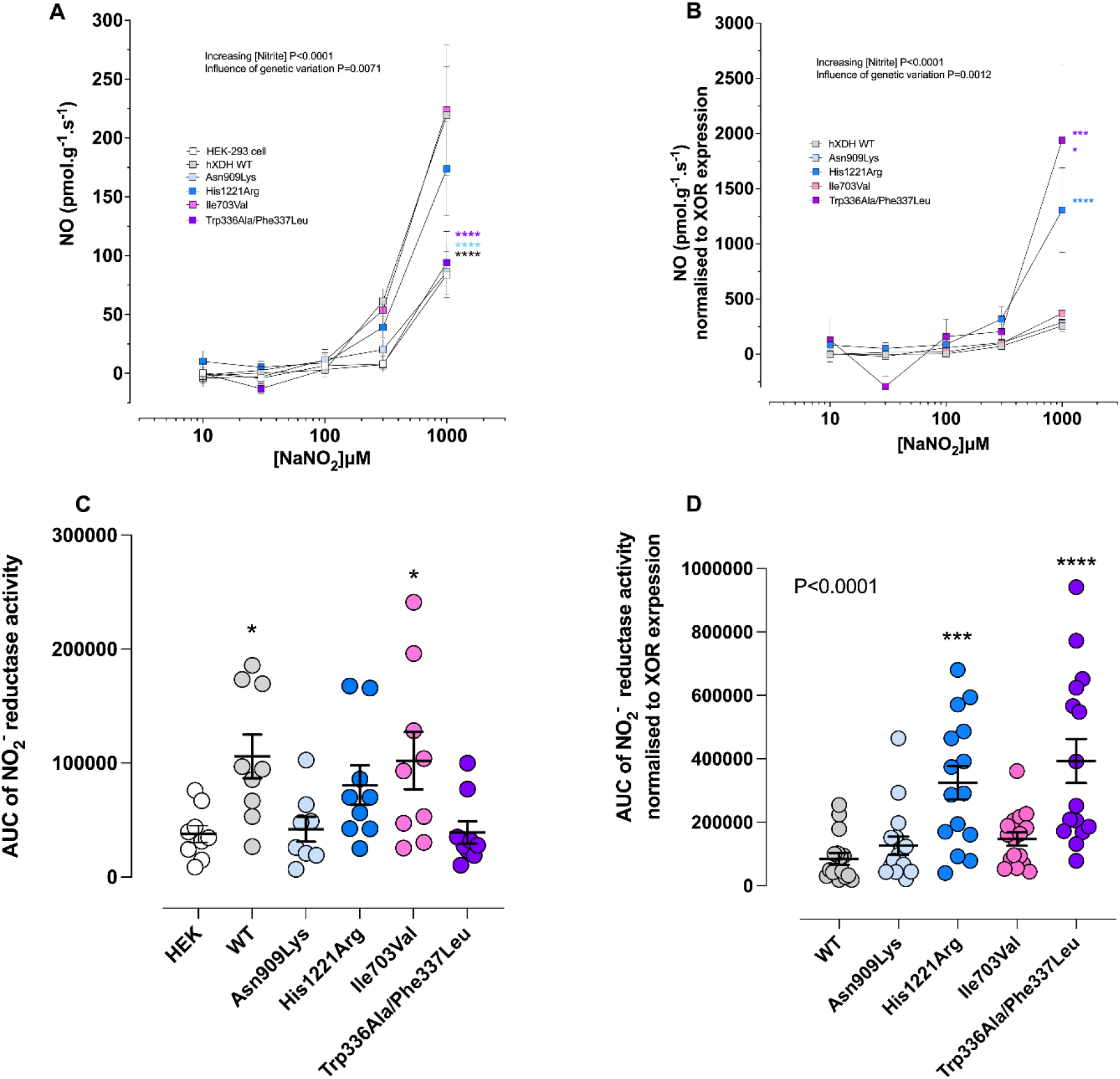
The effects of genetic variation on the NO_2_^-^-reductase activity of XDH. Panel (A) shows the NO_2_^-^-reductase activity measured using ozone chemiluminescence with 250µg of cell lysate of untransfected HEK and hXDH WT, Asn909Lys, His1221Arg, Ile703Val, and Trp336Ala/Phe337Leu expressing HEK cells. Panel (B) shows the ·NO generation following normalization to XOR expression levels of the same samples. Panels (C-D) show the AUC of the baseline NO ^-^-reductase activity. Data are shown as mean ± SEM of n=9 independent experiments. Statistical analysis conducted using two way ANOVA comparing all curves to the hXDH WT control with P-values shown for the variables and post-hoc comparisons at each concentration determined using Sidak’s post-hoc test and one way ANOVA with Dunnett’s post- hoc comparisons for comparison to HEK-293 (C) or hXDH WT (D) and shown as * for P<0.05, ***for P<0.001 and **** for P<0.0001.

**Figure 7.**
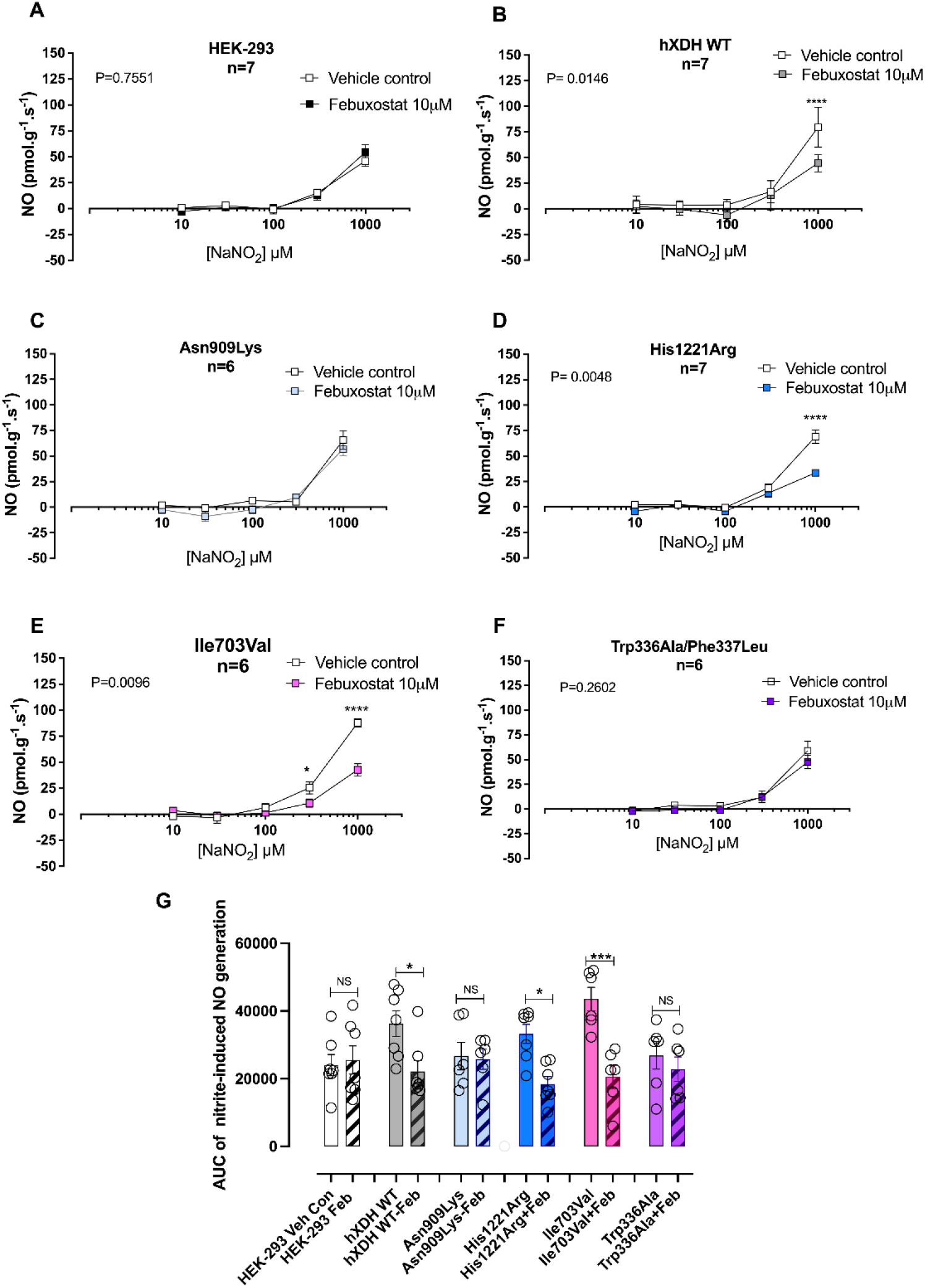
The XOR inhibitor Febuxostat (10µM) prevents NO_2_^-^–reductase activity. Figure A-F show the nitrite-reductase activity in presence of a Mo-Pt specific inhibitor, febuxostat (10µM), and G) shows the AUC of NO_2_^-^-reduction ·NO generation. Statistical analysis conducted using two way ANOVA comparing vehicle vs febuxostat 10µM with P-values shown for the variables and post-hoc comparisons at each concentration determined using Sidak’s post-hoc test (A-F) or one way ANOVA followed by Sidak’s post-hoc test for multiple comparison test (G) and shown as *for P<0.05, ***P<0.005, and **** for P<0.0001.

### Xanthine and NADH increase NO_2_^-^-reductase activity

Since XOR catalyses the one electron reduction of NO_2_^-^ to ·NO in the presence of either xanthine or NADH (37) we measured ·NO formation in the presence of optimal concentrations of each substrate. Interestingly, only the WT and those cells expressing the Ile703Val mutation demonstrated an increase in activity versus the vehicle control in the presence of either substrate (Figure 8). Interestingly, whilst xanthine increased NO_2_^-^ reduction in cells expressing the His1221Arg mutation, no statistically significant effect of NADH was demonstrated (Figure 8G and H).

**Figure 8.**
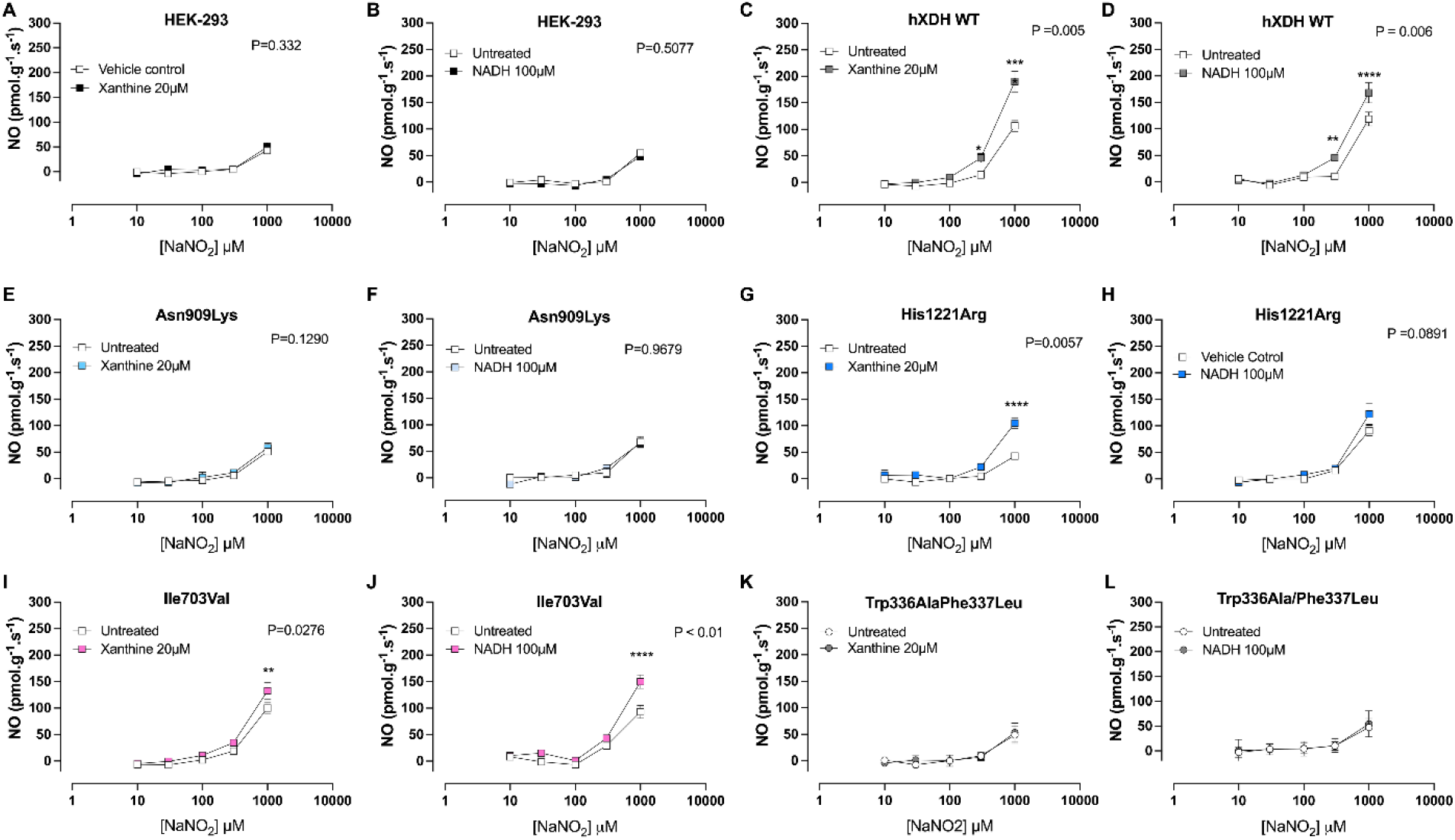
Xanthine and NADH increase NO_2_^-^-reductase in hXDH wild type, His1221Arg and Ile703Val expressing cells. The panels show NO_2_^-^-reductase activity measured by ozone chemiluminescence using 250µg of cell lysate following 5 and 3 min incubation with XOR reducing agents, xanthine 20µM or NADH 100µM respectively, in HEK-293 cells (A-B), hXDH WT (C-D), Asn909Lys (E-F), His1221Arg (G-H), Ile703Val (I-J) and Trp336Ala/Phe337Leu (K-L). Data are shown as mean ± SEM of n=6-8. Statistical analysis conducted using two way ANOVA with Sidak’s post-hoc comparisons between the treatments and shown as **P<0.01 and **** as P<0.0001.

## Discussion

There has been considerable interest in identifying genetic variants of *XDH* that might associate with disease. This is particularly due to the strong associations of disease outcomes with products of XOR activity, most commonly UA for gout and CVD; the latter including hypertension and coronary artery disease. It is notable however, that of the known mutations that exist for the h*XDH* gene, the majority are rare and have been functionally associated with the equally uncommon phenotype of xanthinuria (17,24,38–40). In addition, there is a growing equipoise regarding the clinical utility of targeting the enzyme for the treatment of disease beyond gout (8). There are concerns regarding the chronic use and up titration of inhibitors of the enzyme (41), related to CVD, that has led to questions over the biology of the enzyme. Although a recent Medicines and Healthcare products Regulatory Agency (MHRA) mandated trial has not supported this (42). Surprisingly, clinical trials testing XOR inhibitors in CVD, such as heart failure, have not been positive and in some instances appeared to worsen outcome (43–45). It is thought that at least some of this ambiguity may be due to differences in the extent of XOR upregulation in disease. However, we recently speculated that the current equipoise relates to lack of consideration of XOR’s third biochemical function as a NO_2_ ^-^ reductase (29). For this reason, we re-assessed the biochemical activity of some well-described mutations from the literature that have been associated with xanthinuria or increased UA synthesis and combined this with assessment of the previously unconsidered NO_2_ ^-^-reductase activity of the variant. Herein, we show that previously identified SNPs may have an impact on mRNA stability with direct consequences on XOR expression levels and activity. We also show that non-synonymous substitutions within the peptide chain affect the XDH/XO proportion and consequently its oxidase activity as well as its affinity for the final electron acceptor, NAD^+^ or O_2_, with direct consequences for UA, ROS production and NO_2_^-^-reductase activity. Our findings demonstrate important unexpected impacts upon NO_2_^-^-reductase activity of mutations known to occur in the population that theoretically might improve the redox balance in vivo and so potentially health status. Considering these observations, we suggest that reassessment of both common and rare SNPs in XOR should be explored to include determination of the impact upon NO_2_ ^-^-reductase activity.

XOR is considered to be ubiquitously distributed (46), with the liver and intestine the primary sources, with a subcellular localisation predominantly within the cytoplasm (47) but also on the outer surface of the cell membrane (48). Rouquette et al. also demonstrated a polarised distribution on the cell surface in regions of close cell-to-cell contact. Our confocal studies suggest that the subcellular distribution of XOR, expressed in HEK cells, is cytoplasmic. It is noteworthy that this distribution was largely duplicated by each of the mutants further supporting the validity of our approach for comparing the hXDH WT to the individual mutants.

We observed substantial differences in mRNA expression levels of the mutations versus the h*XDH* WT that was not directly matched to protein expression. This observation is in line with previous findings demonstrating that exonic point mutations may influence mRNA levels by altering its stability (49). It is also possible that the observed changes in gene expression result from variations in the integration process of the external gene within the host genome. However, this is unlikely since the protocols used for each mutation and the host cell source for the expression were identical. The disconnect between mRNA and protein expression was most pronounced for the two mutations Trp336Ala/Phe337Leu and Asn909Lys. Importantly, whilst the former exhibited a diminished level of XOR protein compared to the detected mRNA, suggesting a negative impact of the double mutations on the maturation or stability of the protein, the opposite was observed with the latter. In keeping with our work, a similar study using COS-7 cells, transiently transfected with human *XDH*, showed differential protein expression levels across the investigated mutants (22). Interestingly, in this study in COS-7 cells the Mo-Co mutation Asn909Lys, almost eliminated detectable protein. This contrasting observation may relate to the experimental approach used or it may suggest a cell-specific expression effect. Indeed, in support of this observation a study published by Kudo et al.(28) demonstrated that the luciferase reporter assay, used to evaluate whether mutations within the promoter region of *XDH* have an effect upon XOR expression, revealed significant differences between investigated SNPs as well as different cell lines.

Numerous pathological stimuli including pro-inflammatory cytokines (50–52), unhealthy diet (53), cigarette smoke (13), hypoxia (14,54), ischemia-reperfusion injury and CVD risk factors (such as hypertension (55)) increase XOR expression and activity. These stimuli also promote the conversion of XOR from the housekeeping reduced isoform, XDH, to the more detrimental oxidized isoform, XO. Because of this modification, the XDH/XO ratio shifts in favour of oxygen radical production. Our study demonstrates that the SNPs leading to non-synonymous substitutions have a critical role in determining XDH/XO proportion. We observed that His1221Arg and Ile703Val exhibited a prevalence of the oxidase isoform of the enzyme over the dehydrogenase. It is possible that this observation is due to the conversion of the NAD^+^- dependency to O_2_-dependency as a consequence of sample preparation (1) or due to low temperature storage of the cell extract (56), although since all samples were treated identically we think this is unlikely. The His1221Arg mutant exhibited greater XO activity of 70% of the total compared to the WT (55%) and Ile703Val (57%). Therefore, we propose that the dehydrogenase conversion to the oxidase isoform is not only an effect of post-translational modification but may also be a consequence of non-synonymous alteration of the peptide chain. Nishino et al., using a non-mammalian expression system, demonstrated that non-synonymous substitutions of key cysteine residues (Cys535Ala and Cys992Arg) prevent XDH oxidation (57).

Importantly, we acknowledge that of the mutations we tested only the artificial Trp336Ala variant resides with the active site, however, our observations indicate that mutations distant from this site also impact biochemical activity. Interestingly, both Cys535 and 992 are well conserved across mammalian XDHs indicating the generalisability of this regulation across species. The one exception is chicken where the electrically neutral Cys992 is replaced by the positively charged Arg (58). Furthermore, it is noteworthy that, whilst chicken Cys535 is followed by glutamic acid (Glu) at the position 536, mammalian Cys535 is followed by glycine (Gly). This difference, whilst seemingly unimportant, prevents the formation of a disulphide bridge leading to speculation that other adjacent amino acids may be critical in determining conversion (59). This leads us to speculate that His1221 may still influence Cys(s) oxidation capacity and consequently XDH conversion to XO. Indeed, X-ray crystallography of purified bovine XOR showed that, when completely folded, regions considered spatially distant, i.e. the Mo-Pt domain and the interfaces of the Fe/S and FAD domains, come into close contact (60). In keeping with these observations, our computational analysis reveals that His1221 interacts with the backbone and sidechain of Trp1329 likely stabilising the loop structures in this region. Thus, the change to Arg may impact upon the oxidation process of adjacent amino acids, including Cys1326 which has been also implicated in conversion of XDH to XO (57). However, regardless of the nature of the process, reversible or irreversible, XDH conversion to XO is accompanied by a disruption of a central cluster (Arg335, Trp336, Arg427, and Phe550) with consequent conformational changes and relocation of the active loop A around the FAD site (57). There is evidence that a double mutation Trp336Ala/Phe337Leu, also reproduced in our study, is accompanied by a loss of interaction of Phe549 (human 550) favouring XO over the XDH structure (30).

Whilst most mutations identified in the *XDH* open reading frame have been associated with a reduced or abolished XOR activity (17), a few have been associated with an increased oxidase activity (19,61). Of the 21 non-synonymous mutations previously identified, 10 exhibited significant differences in the kinetic parameters (Km, Vmax, Vmax/Km) of XOR-driven xanthine oxidation. Amongst these, only two, His1221Arg and Ile703Val, showed a significant increase in purinergic activity compared to the WT due to a substantial rise of Vmax (22), whereas Asn909Lys demonstrated a significant loss of activity. In agreement, we confirmed that non-synonymous mutations of the C-terminal domain, His1221Arg and Ile703Val, modulate the oxidase activity of the enzyme which may account for inter-individual variations of oxidative stress (19,20,61). However, in contrast to Kudo et al., we observed a significant increase in O_2_^·-^ production with His1221Arg in presence of both reducing agents, xanthine and NADH, and a similar activity with xanthine for Ile703Val versus the WT. A plausible rationale for this discrepancy may be that in the Kudo et al study, UA, a surrogate of XOR activity, was the only sub-product measured and the obtained values were not normalized to the enzyme expression levels. This explanation is further corroborated by our UA measurement which reveals important differences between pre- and post-normalization to XOR expression levels with His1221Arg causing statistically significant elevation of UA levels at baseline.

Our findings regarding Asn909Lys activity also show differences to those previously reported (22,30). Indeed, whilst Kudo et al. demonstrated a 4-fold reduction of Vmax compared to WT for UA synthesis highlighting the mutation as a potential cause of xanthinuria, our O_2_^·-^ analysis only revealed a downward trend compared to hXDH WT, whereas UA quantification showed similar activity in presence of xanthine. In addition, when the FAD-reducing agent NADH, was provided, and the activity was normalised to XOR expression the Asn909Lys showed similar levels of O_2_^·-^ to hXDH WT, suggesting that the reductions are likely due to lower levels of expression rather than lower activity per se. Similarly, the artificial double mutant, Trp335Ala/Phe336Leu (human Trp336Ala/Phe337Ala), suggested to be a highly O_2_^·-^-producing enzyme (30), exhibited a profoundly reduced xanthine-driven dioxygen reduction capacity and no difference with NADH. However, when normalised to XOR expression level in the presence of NADH this mutant expressed a far greater production of O_2_^·-^ than WT or His1221Arg; again suggesting that the much lower expression level of the enzyme influenced perceived changes in activity. It has been suggested that His1221Arg is localized at the surface of the Mo-Pt binding domain and, even though not directly involved in the formation of the active site cavity (Glu803, Arg881, Glu1262), it positively modulates UA production by increasing its release rate (23). There is also some evidence suggesting that chemical modification of surface amino acid residues slow XOR activity with a consequent reduction in urate release (62). Taken together, these observations confirm the direct involvement of previously thought non-critical amino acid residues in the oxidation process of purines to UA, offering a plausible explanation for the increased O_2_ ^·-^ production observed with His1221Arg. Our observations are important since they clarify better whether the mutations alter activity or simply expression level of the protein. However, we suggest that the increase in dioxygen reduction observed in this study, results from a combination of increased oxidase activity, as observed by Kudo et al.(28), along with altered XDH/XO proportion, making His1221Arg a potential marker of hyperuricemia, oxidative stress, and endothelial dysfunction.

As mentioned the functional remit of XOR has extended well beyond ROS and UA synthesis over the past 10-15 years, with the growing appreciation that in disease, particularly demonstrated for CVD, XOR acts as a critical NO_2_^-^-reductase (63–65). Considering these observations, the increase in NO_2_^-^ reduction to ·NO observed with some of the naturally occurring mutations may be particularly important for health. NO_2_^-^-reductase activity for His1221Arg and Trp336Ala/Phe337Leu was elevated in comparison to WT with a trend for Ile703Val. The possibly improved ·NO formation observed with Ile703Val compared to hXDH WT may be of particular importance considering that this mutant exhibited a similar oxidase activity in presence of both reducing agents. This finding opens a new important question about the potential protective role of Ile703Val and His1221Arg which warrants further structural and molecular investigations. On the contrary, the increased NO_2_ ^-^-reductase activity of the artificial double mutant appears to be less relevant as the XOR-driven NO_2_ ^-^ reduction was not attenuated by XOR inhibition. Previous in vitro and in vivo observations have shown that the beneficial effects resulting from the non-canonical pathway of ·NO generation in experimental models and patients with CVD are absent following XOR inhibition (6,55).

To interrogate more closely the possibility that non-synonymous mutations may alter the enzymatic tertiary structure and consequently the intramolecular shuttling of electrons, we investigated the NO_2_^-^-reductase activity in the presence of NADH, or xanthine. These substrates did not influence ·NO generation from NO_2_^-^ in HEK-293, Asn909Lys or Trp336Ala/Phe337Leu cell lysate. In contrast for cells expressing hXDH WT, His1221Arg or Ile703Val, NO_2_ ^-^ reduction was facilitated in the presence of both reducing agents, with His1221Arg showing a preference for xanthine over NADH. Taken together these findings suggest that in His1221Arg, the two redox centres Fe/S I and II are still functional and efficiently shuttling e^-^ between the Mo-Pt and FAD sites; both xanthine- and NADH-derived e^-^ are able to catalyse dioxygen and NO_2_^-^ reductions; and finally, the e^-^ passed to the enzyme are preferentially used at the same site of production and thus His1221Arg showing a tendency of more efficiently using xanthine over NADH and vice versa for Ile703Val. These observations, albeit exploratory, suggest that non-synonymous mutations of the enzyme may change its affinity for Mo-Pt or FAD specific substrate. Considering that purines and NADH concentrations change dramatically before and after a pathological insult, such as ischemia, this aspect warrants further investigations.

We acknowledge that our study utilises an in vitro expression system and thus whilst the cell system is of human origin further in vivo analysis of the consequences of these mutations is needed. This can be addressed by creation of transgenic mouse models as well as greater scrutiny of large genome-wide-association-study (GWAS) for further confirmation through replication. Since many known mutations of XDH are rare, recent advances in testing the impact of rare mutations upon key disease traits and markers is warranted.

## Conclusion

In this study we have shown that a naturally occurring non-synonymous mutation of human XDH, His1221Arg leads to an altered XDH/XO proportion and an increased oxidase activity; effects typically seen in numerous pathological conditions. However, we have also demonstrated that the upregulation of XOR activity is accompanied by a substantial rise in its NO_2_^-^-reductase potential. We also suggest that specific mutations such as Ile703Val, with a similar oxidase activity of WT but potentially increased NO_2_^-^-reductase activity, may exert a biologically important effect in preventing aspects of CVD (Table 2). Collectively, these data lead us to speculate that carriers of a potential detrimental mutation, His1221Arg, may revert their latent pathological phenotype to a more beneficial one by targeting the non-canonical pathway for ·NO generation through the safe application of dietary inorganic nitrate (NO_3_^-^). With emerging evidence suggesting elevated sUA, ROS, and ·NO depletion are key markers of CVD, identification of pro-oxidative and uricaemic genotypes would allow potential stratification to treatments that might be useful in elevating NO levels leading to health benefits.

**Table 2:** Summary table indicating fold changes of hXDH mutants relative to XDH WT.

| Mutation | Gene and protein expression |  | XDH and XO pterin proportions |  | O<sub>2</sub><sup>-</sup> generation |  | UA | NO<sub>2</sub><sup>-</sup> reductase activity |
| --- | --- | --- | --- | --- | --- | --- | --- | --- |
|  | XDH mRNA | XDH protein | XDH | XO | Xanthine | NADH |  |  |
| hXDH WT | 1 ± 0.2 | 1 ± 0.1 | 1 ± 0.1 | 1 ± 0.1 | 1 ± 0.5 | 1 ± 0.2 | 1 ± 0.3 | 1 ± 0.9 |
| Asn909Lys | 0.2 ± 0.03 | 0.4 ± 0.1 | 0 ± 0 | 0 ± 0 | 0.01 ± 0.02 | 1.2 ± 0.7 | 0.9 ± 0.3 | 1.5 ± 1.3 |
| His1221Arg | 0.3 ± 0.08 | 0.2 ± 0.03 | 0.7 ± 0.2 | 1.2 ± 0.1 | 4.1 ± 1.7 | 4.2 ± 1.5 | 6.6 ± 3.9 | 3.8 ± 2.4 |
| Ile703Val | 0.4 ± 0.09 | 0.7 ± 0.07 | 0.9 ± 0.2 | 1.0 ± 0.2 | 1.1 ± 0.6 | 1.3 ± 0.3 | 1.9 ± 0.9 | 1.7 ± 1.0 |
| Trp336Ala | 0.5 ± 0.07 | 0.1 ± 0.04 | 0 ± 0 | 0 ± 0 | 0 ± 0 | 6.4 ± 3.1 | 3.3 ± 1.6 | 4.6 ± 3.2 |

## This article contains supplementary materials Acknowledgments

### General

We would like to express our gratitude to Professor Richard M. Wright for providing the pcDNA.3.1-myc-hysA-hXDH plasmid vector. We also wish to thank Dr Claudio Raimondi for assisting in acquiring images via confocal microscopy.

### Funding

GM was partially supported by Menarini Ricerche (awarded to Prof Claudio Borghi) and the Italian society of Pharmacology (SIF). This work and GM were also supported by The Barts Charity Cardiovascular Programme (MRG00913). Nicki Dyson is funded by a BHF MRes/PhD Studentship (FS/19/62/34901).

**Abbreviations**
AUC: Area Under the Curve
Asn909Lys: Asparagine909Lysine
CVD: Cardiovascular disease
CPS: Count per second
Cys: Cysteine
DPI: Diphenyleneiodonium
FAD: Flavin adenine dinucleotide
GWAS: Genome-wide-association study
His1221Arg: Histidine1221Arginin
HEK-293: Human Embryonic Kidney
hXDH WT: Human Xanthine Dehydrogenase wild-type
H_2_O_2_: Hydrogen peroxide
Fe-S: Iron-sulphur
Ile703Val: Isoleucine703Valine
LECL: Lucigenin-enhanced chemiluminescence
MHRA: Medicines and Healthcare products Regulatory Agency
Mo-Pt: Molybdo-Pterine
NAD^+^: Nicotinamide adenine dinucleotide
NADH: Nicotinamide adenine dinucleotide hydrogen
NO^-^_3_: Nitrate
NO^-^_2_: Nitrite
·NO: Nitric oxide
O_2_: Oxygen
ROS: Reactive oxygen species
Trp336Ala/Phe337Leu or Trp336Ala: Tryptophan336Alanine/Phenylalanine337Leucine
SNPs: Single nucleotide polymorphisms
O_2_^·-^: Superoxide anion
UA: Uric acid
XDH: Xanthine dehydrogenase
XOR: Xanthine Oxidoreductase
XO: Xanthine Oxidase

