## supplement material for "Natural mutations of human *XDH* promote the nitrite (NO_2_^-^)-reductase capacity of xanthine oxidoreductase: a novel mechanism to promote redox health?"

**List of material Included**

**1. Table S1**
**2. Table S2**
**3. Table S3**
**4. Table S4**
**5. Supplement Figure 1 (S1)**
**6. Supplement Figure 2 (S2)**

**Table S1 Primers used for Site Directed Mutagenesis of hXDH WT.** Table S1 shows the primers, forward and reverse, used to generate each mutant. All the mutagenic primers were designed using the NEBaseChanger software (http://nebasechanger.neb.com).

| Mutation | Nucleotide substitution | Forward primer | Reverse primer | Annealing Temperature (Ta) |
|---|---|---|---|---|
| Asn909Lys | 2727 C > A | TTCCCTCCAAaACGGCCTTCC | GGTTGGTTTTGCACAGCC | 66 °C |
| His1221Arg | 3662 A > G | GGGAGCCTGCgCACCCGTGGC | CTCGGGGGAATAGTGTAGCTCCTCTAGG | 72 °C |
| Ile703Val | 2107 A > G | TGAGGATGCTgTAAAGAACAAC | ATTGTGATAATGGCTGGTAG | 59 °C |
| Trp336Ala/ Phe337Leu | 1006/7 TG > GC 1009 T > C | GCAGCTGCGCgcGTTTGCTGGG GCTGCGCTGGcTTGCTGGGAA | TCCAGGACCCCTCTGAAC TGCTCCAGGACCCCTCTG | 66 °C 70 °C |

**Table S2 PCR cycling condition used for Side Directed Mutagenesis.** Table S2 shows the PCR cycling conditions used for Site directed mutagenesis. All the conditions are according to the manual except the annealing temperature, Ta, which varied according to the primers used, and was automatically generated by the NEB primer design software (NEBaseChanger.neb.com).

| Step | Temperature | Time |
|---|---|---|
| Initial denaturation | 98 °C | 30 seconds |
| 25 cycles | 98 °C | 10 seconds |
| | Ta (see Table S1) | 30 seconds |
| | 72 °C | 4 minutes |
| Final extension | 72 °C | 2 minutes |
| Hold | 4 °C | |

**Table S3 Primers used for sequencing.** Table shows the primers used for the Sanger Sequencing of hXDH WT and mutant forms. Each primer covered roughly 300bp. Two reverse primers were used to ensure that the 5' region of the cDNA was completely covered.

| **Primer (5' 3')** | **Length** | **% GC** |
|---|---|---|
| 1st Forward Primer **AAGACGAGGCTGCATCCTGT** | 20bp | 55 |
| 2nd Forward Primer **GGGAACACGGAGATTGGCAT** | 20bp | 55 |
| 3rd Forward Primer **AGGTAACCAGTGGCATGAGA** | 20bp | 50 |
| 4th Forward Primer **CGCTACGAGAATGAGCTGTC** | 20bp | 55 |
| 5th Forward Primer **GGTGGCCAAGAGCACTTCTA** | 20bp | 55 |
| 6th Forward Primer **ACCAACCTTCCCTCCAACAC** | 20bp | 55 |
| 7th Forward Primer **AACACTGTGCCCAACACCTC** | 20bp | 55 |
| 8th Forward Primer **CATTGAGTTCAGGGTGTCCCT** | 20bp | 55 |
| REV/Complement primer **ACAGGATGCAGCCTCGTCTT** | 20bp | 55 |
| REV/COMP_Seq_2 **CTCCACAGCATCCACCATC** | 19bp | 58 |

**Table S4 Primers used for qPCR**. Table S4 shows the sequence of the primers, forward and reverse, used for the qPCR analysis of hXDH, β-actin and GAPDH genes.

| Gene | Forward primer (5' ⟶ 3') | Reverse primer (5' ⟶ 3') |
|---|---|---|
| Human XDH | CTCAGTCAGCCTCTCGCCAT | TATCCACGTCACACGCTCCC |
| Human β-actin | TCGTGCGTGACATTAAGGAG | AGGAAGGAAGGCTGGAAGAG |
| Human GAPDH | TGGTCTCCTCTGACTTCAAC | GTGAGGGTCTCTCTCTTCCT |

**Supplement Figure 1 (S1). Figure 3 Intracellular XOR expression detected via flow cytometry.** Figure A) shows representative histograms of permeabilised cells showing XOR expression of WT and mutants. Red histogram depicts cells incubated with secondary antibody alone as a control whilst the blue histogram depicts cells incubated with primary XOR and secondary antibody. Figure also shows XOR quantification as percentage of cells expression B) and as percentage of increase in median florescence intensity (MFI) compared to secondary alone C). All data are expressed as mean ± SEM of n=5 independent experiments and compared using one-way ANOVA with Dunnett post hoc analysis comparing to the hXDH WT control and post-test significance shown as *P<0.05, P<0.01, P<0.0001.

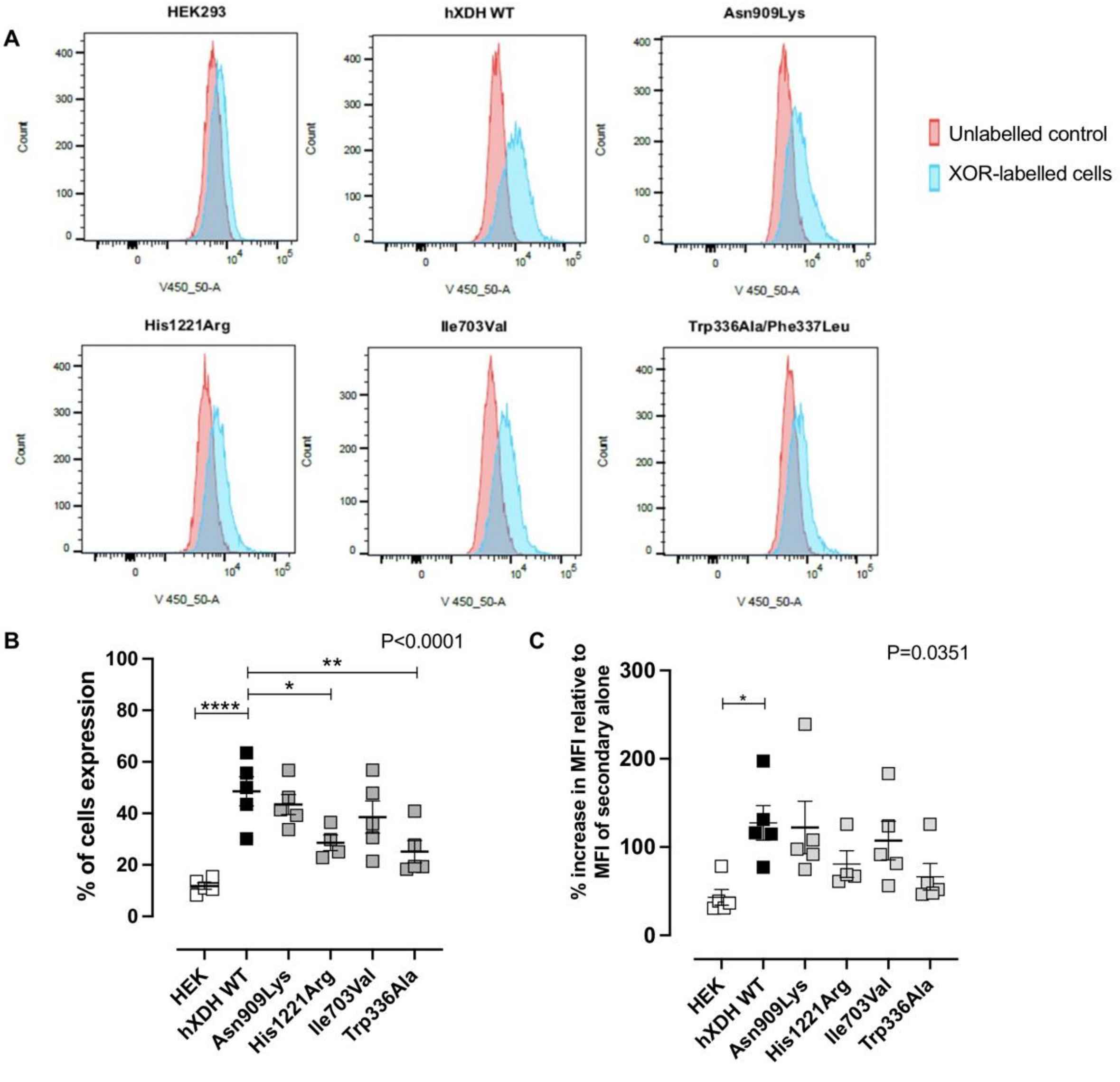

**Supplement Figure 2 (S2). Immunoblotting of XOR and GAPDH in increasing concentration of stable hXDH WT cell homogenate.** Figure A and B show XOR and GAPDH expression with increasing amounts (1, 5, 10, 50, 100, 250µg) of cell homogenate. Both XOR and GAPDH (housekeeping protein used to normalise XOR expression) were within the linear range when loading 50µg of cell lysate. Figure C and D are representative immunoblots of XOR and GAPDH. Data are shown as SEM of n=3 independent experiments.

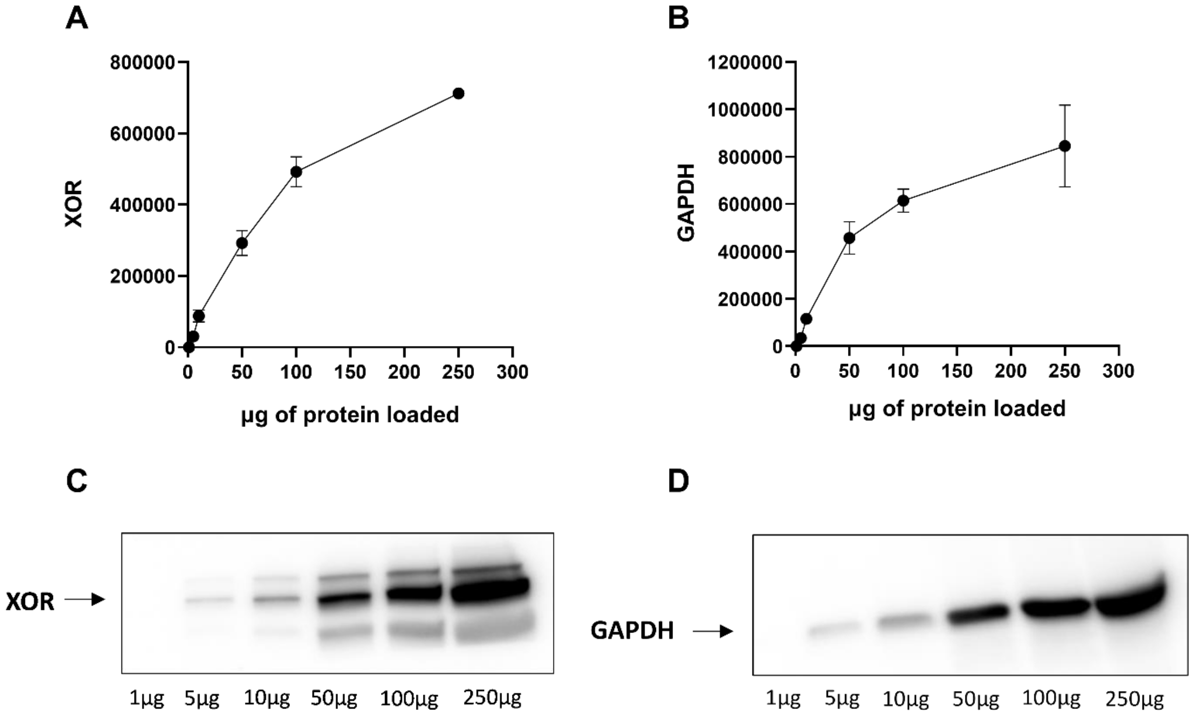
